# PyRates – A Python Framework for rate-based neural Simulations

**DOI:** 10.1101/608067

**Authors:** Richard Gast, Daniel Rose, Harald E. Möller, Nikolaus Weiskopf, Thomas R. Knösche

## Abstract

In neuroscience, computational modeling has become an important source of insight into brain states and dynamics. A basic requirement for computational modeling studies is the availability of efficient software for setting up models and performing numerical simulations. While many such tools exist for different families of neural models, there is a lack of tools allowing for both a generic model definition and efficiently parallelized simulations. In this work, we present PyRates, a Python framework that provides the means to build a large variety of neural models as a graph. PyRates provides intuitive access to and modification of all mathematical operators in a graph, thus allowing for a highly generic model definition. For computational efficiency and parallelization, the model graph is translated into a *tensorflow*-based compute graph. Using the example of two different neural models belonging to the family of rate-based population models, we explain the mathematical formalism, software structure and user interfaces of PyRates. We then show via numerical simulations that the behavior shown by the model implementations in PyRates is consistent with the literature. Finally, we demonstrate the computational capacities and scalability of PyRates via a number of benchmark simulations of neural networks differing in size and connectivity.

## Introduction

In the last decades, computational neuroscience has become an integral part of neuroscientific research. A major factor in this development has been the difficulty to gain mechanistic insights into neural processes and structures from recordings of brain activity without additional computational models. This problem is strongly connected to the signals recorded with non-invasive brain imaging techniques such as magneto- and electroencephalography (MEG/EEG) or functional magnetic resonance imaging (fMRI). Even though the spatiotemporal resolution of these techniques has improved throughout the years, they are still limited with respect to the state variables of the brain they can pick up. Spatial resolution in fMRI has been pushed to the sub-millimeter range [1, 2], whereas EEG and MEG offer a temporal resolution thought to be sufficient to capture all major signaling processes in the brain [3]. On the EEG/MEG side, the measured signal is thought to arise mainly from the superposition of primary and secondary currents resulting from post-synaptic polarization of a large number of cells with similarly oriented dendrites [4]. Therefore, the activity of cell-types that do not show a clear orientation preference (like most inhibitory interneurons [5]) can barely be picked up, even though they might play a crucial role for the underlying neural dynamics. Further issues of EEG/MEG acquisitions are their limited sensitivity to sub-cortical signal sources and the inverse problem one is facing when trying to locate the source of a signal within the brain [6]. On the other hand, fMRI measures hemodynamic signals of the brain related to local blood flow, blood volume and blood oxygenation levels and thus delivers only an indirect, strongly blurred view on the dynamic state of the brain [7]. These limitations pose the need for additional models and assumptions that link the recorded signals to the underlying neural activity. Computational models of brain dynamics (called neural models henceforth) are therefore particularly important for interpreting neuroimaging data and understanding the neural mechanisms involved in their generation [8–10]. Such models have been developed for various spatial and temporal scales of the brain, ranging from highly detailed models of a single neuron to models that represent the lumped activity of thousands of neurons. In any case, they provide observation and control over all state variables included in a given model, thus offering mechanistic insights into their dynamics.

Numerical simulations are the major method used to investigate neural models beyond pure mathematical analyses and link model variables to experimental data. Such numerical simulations can be computationally highly expensive and scale with the model size, simulation time and temporal resolution of the simulation. Different software tools have been developed for neural modeling that offer various solutions to render numerical simulations more efficient (e.g. TVB [11], DCM [12], Nengo [13], NEST [14], ANNarchy [15], Brian [16], NEURON [17]). Since the brain is an inherently highly parallelized information processing system (i.e. all of its 10 billion neurons transform and propagate signals in parallel), most models of the brain have a high degree of structural parallelism as well. This means that they involve calculations that can be evaluated in parallel, as for example the update of the firing rate of each cell population inside a neural model. One obvious way of optimizing numerical simulations of neural models is therefore to distribute these calculations on the parallel hardware of a computer, i.e. its central and graphical processing units (CPUs and GPUs). Neural simulation tools that implement such mechanisms include Nengo [13], ANNarchy [15], Brian [16], NEURON [18] and PCSIM [19], for example. Each of these tools has been build for neural models of a certain family. While complex multi-compartment models of single spiking neurons are implemented in NEURON, models of point neurons are provided by ANNarchy, Brain, Nengo, Nest and PCSIM, whereas neural population models are found in TVB and DCM. For most of these tools, a pool of pre-implemented models of the given family are available that the user can choose from. However, most often it is not possible to add new models or modeling mechanisms to this pool without considerable effort. This holds true especially, if one still wants to benefit from the parallelization and optimization features of the respective software. Exceptions are tools like ANNarchy and Brian that include code generation mechanisms. These allow the user to define the mathematical equations that certain parts of the model will be governed by and will automatically translate them into the same representations that the pre-implemented models follow. Unfortunately, the tools that provide such code generation mechanisms are limited with regards to the model parts they allow to be customized in such a way and the families of neural models they can express.

To summarize, we believe that the increasing number of computational models and numerical simulations in neuroscientific research motivates the development of neural simulation tools that:

- follow a well-defined mathematical formalism in their model configurations,
- are flexible enough so scientists can implement custom models that go beyond pre-implemented models in both the mathematical equations and network structure,
- are structured in a way such that models are easily understood, set up and shared with other scientists,
- enable efficient numerical simulations on parallel computing hardware.

In this work, we present PyRates, a Python framework which is in line with these suggestions (available from https://github.com/pyrates-neuroscience/PyRates). The basic idea behind PyRates is to provide a well documented, thoroughly tested and computationally powerful framework for neural modeling and simulations. Thereby, our solution to the parallelization issue is to translate every model implemented in PyRates into a tensorflow [20] graph, a powerful compute engine that provides efficient CPU and GPU parallelization. Each model in PyRates is represented by a graph of nodes and edges, with the former representing the model units (i.e. single cells, cell populations,…) and the latter the information transfer between them. As we will explain in more detail later on, the user has full control over the mathematical equations that nodes and edges are defined by. Still, both the model configuration and simulation can be done within a few lines of code. In principle, this allows to implement any kind of dynamic neural system that can be expressed as a graph. However, for the remainder of this article, we will focus on a specific family of neural models, namely rate-based population models (hence the name PyRates), which will be introduced in the next section. The focus on population models is (i) in accordance with the expertise of the authors and (ii) serves the purpose of keeping the article concise. However, even though neural population models were chosen as exemplary models, the emphasize of the paper lies on introducing the features and capacities of the framework, how to define a model in PyRates and how to use the software to perform and analyze neural simulations. Therefore, we first introduce the mathematical syntax used for all our models, followed by an explanation how single mathematical equations are structured in PyRates to form a neural network model. To this end, we provide a step-by-step example of how to configure and simulate a particular neural population model. We continue with a section dedicated to the evaluation of different numerical simulation scenarios. First, we validate the implementation of two exemplary neural population models in PyRates by replicating key behaviors of the models reported in their original publications. Second, we demonstrate the computational efficiency and scalability of PyRates via a number of benchmarks. These benchmarks constitute different realistic test networks that differ in size and sparseness of their connectivity. Furthermore, we discuss the strengths and limitations of PyRates for developing and simulating neural models.

## Neural Population Models

Investigating the human brain via EEG/MEG or fMRI means working with signals that are assumed to represent changes in the average activity of large cell populations. While these signals could in theory be explained by detailed models of single cell processes, such models come with a state space of much higher dimensionality than the measured signals and would thus need to be reduced again for comparison. Additionally, the interpretability of single cell activities would be limited, since an average signal is modelled that could result from endlessly many different single cell activation patterns. As an alternative, neural population models (also called neural mass models) have widely been used [21]. They are biophysically motivated non-linear models of neural dynamics that describe the average activity of large cell populations in the brain via a mean-field approach [22–24]. Often, they express the state of each neural population by an average membrane potential and an average firing rate. This allows for a more direct comparison to EEG/MEG and fMRI signals, since the changes in the average activity across many cells are thought to generate those signals [25, 26]. The dynamics and transformations of these state variables can typically be formulated via three mathematical operators. The first two describe the input-output structure of a single population: While the rate-to-potential operator (RPO) transforms synaptic inputs into average membrane potential changes, the potential-to-rate operator (PRO) transforms the average membrane potential into an average firing rate output. Widely used forms for these operators are a convolution operation with an alpha kernel for the RPO (e.g. [24, 25, 27]) and a sigmoidal, instantaneous transformation for the PRO (e.g. [23, 28, 29]). The third operator is the coupling operator (CO) that transforms outgoing into incoming firing rates and is thus used to establish connections across populations. By describing the dynamics of large neural population networks via three basic transforms (RPO, PRO & CO), neural populations combine computational feasibility with biophysical interpretability. Due to these desirable qualities, they have become an attractive method for studying neural dynamics on a meso- and macroscopic scale [8, 10, 21]. They have been established as one of the most popular methods for modeling EEG/MEG and fMRI measurements and were able to account for various dynamic properties of experimentally observed neural activity [26, 27, 30–35].

A particular neural population model we will use repeatedly in later sections is the three-population circuit introduced by Jansen and Rit [24]. The Jansen-Rit circuit (JRC) was originally proposed as a mechanistic model of the EEG signal generated by the visual cortex [24, 36]. Historically, however, it has been used as a canonical model of cell population interactions in a cortical column [30, 31, 35]. Its basic structure can be seen in Figure 1 B, which can be thought of as a zoom-in on a single cortical column. The signal generated by this column is the result of the dynamic interactions between a projection cell population (PC), an excitatory interneuron population (EIN) and an inhibitory interneuron population (IIN). For certain parametrizations, the JRC has been shown to be able to produce key features of a typical EEG signal, such as the waxing-and-waning alpha oscillations [24, 25, 37]. A detailed account of the model’s mathematical description will be given in the next section, where we will demonstrate how to implement models in PyRates, using the example of the JRC equations. We chose to employ the JRC as an exemplary population model in this article, since it is an established model used in numerous publications the reader can compare our reports against.

**Fig 1.**
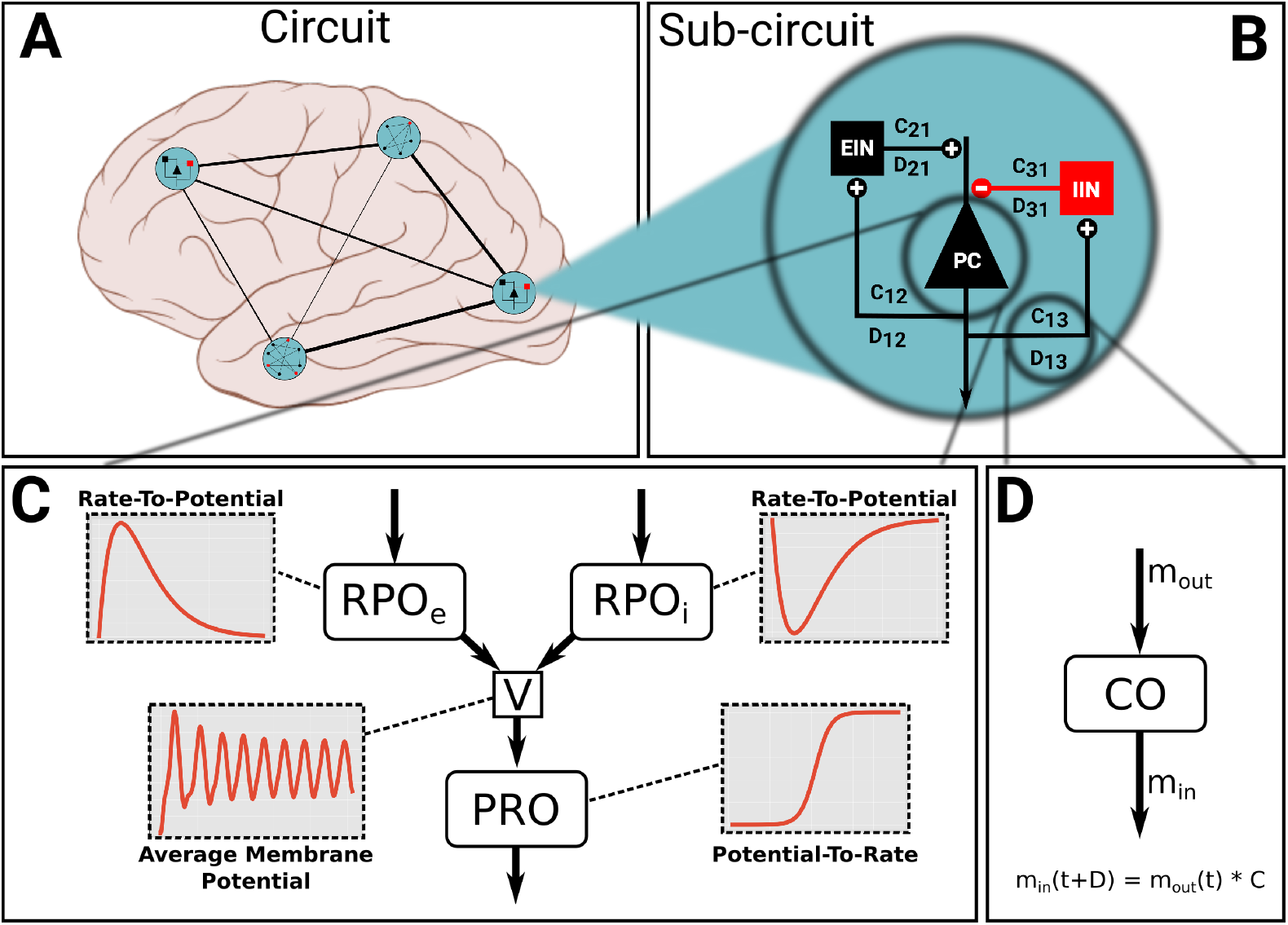
Model structure in PyRates. The largest organisational unit of a network model is the *Circuit*. Any circuit may also consist of multiple hierarchical layers of subcircuits. Panel (A) depicts an imaginary circuit that encompasses four subcircuits that represent one brain region each. One of these local circuits is a Jansen-Rit circuit (B), consisting of three neural populations (PC, EIN, IIN) and connections between them. One node (C) may consist of multiple operators that contain the mathematical equations. Here, two rate-to-potential operators (RPO) convolute incoming firing rates with an alpha kernel to produce post-synaptic potentials. These are summarized int a mean membrane potential *V*. The potential-to-rate operator (PRO) transforms V into outgoing firing rate *m_o_ut* via a sigmoid function. Inset graphs give a qualitative representation of the operators and evolution of the membrane potential. Edges (lines in A and B) represent information transfer between nodes. As panel (D) shows, edges may also contain operators. By default, edges apply a multiplicative weighting constant and can optionally delay the information passage with respect to time. The equation shown in panel (D) depicts this default behaviour.

Another neural population model that we will make use of in this paper is the one described by Montbrió and colleagues [38]. It has been mentioned as one of the next generation neural mass models that provide a more precise mean-field description than classic neural population models like the JRC [39]. The model proposed by Montbrió and colleagues represents a mathematically exact mean-field derivation of a network of globally coupled quadratic integrate-and-fire neurons [38]. It can thus represent every macroscopic state the single cell network may fall into. This distinguishes it from the JRC, since it has no such correspondence between a single cell network and the population descriptions. Furthermore, the macroscopic states (average membrane potential and average firing rate) of the Montbrió model can be linked directly to the synchronicity of the underlying single-cell network, a property which benefits the investigation of EEG phenomena such as event-related (de-)synchronization. We chose this model as our second example case due to its novelty and its potential importance for future neural population studies. Within the domain of rate-based neural population models, we found these two models sufficiently distinct to demonstrate the ability of PyRates to implement different model structures.

## The Framework

PyRates is a framework to construct and simulate computational neural network models. The core goal is to let scientists focus on the model building, *i.e.* defining model structure and working out the equations – while the software takes care of setting up the network, implementing equations and optimizing the computational workload.

This goal is reflected in the modular software design and user interface. Model configuration and simulation are realized as separate software layers as depicted in Figure 2. The frontend features user interfaces for different levels of programming expertise and allows scientists to flexibly implement custom models. These are transformed into a graph-based intermediate representation that the backend interprets to perform efficient computations. We employ a custom mathematical syntax and domain specific model definition language. Both focus on readability and are much reduced in comparison to general-purpose languages. The following paragraphs explain the user interfaces to define models and run simulations, which is the focus of this paper. More details on implementation can be found in the online documentation (see https://github.com/pyrates-neuroscience/PyRates).

**Fig 2.**
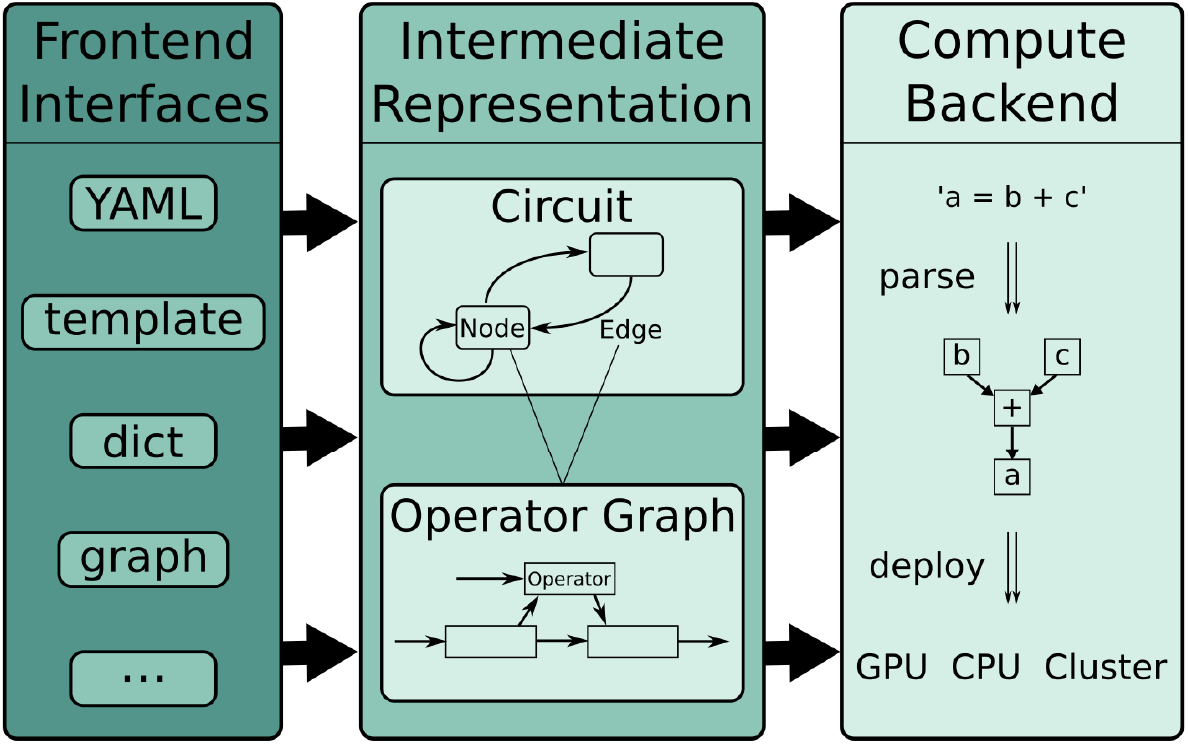
Schematic of software layers. PyRates is separated into frontend, intermediate representation (IR) and backend. The frontend features a set interfaces to define network models. These are then translated into a standardized structure, called the IR. Simulations are realized via the backend, which transforms the high-level IR into lower-level representations for efficient computations. The frontend easily can be extended with new interfaces, while the backend can be swapped out to target a different computation framework.

### Mathematical syntax

Neural network models are usually defined by a set of (differential) equations and corresponding parameters. Scientists can define computational models in terms of algebraic equations and relations between different equations in PyRates. The software interprets and implements these equations prior to a simulation to perform efficient computations. This includes the implementation of numerical integration schemes for 1st order differential equations and parallelization. The mathematical syntax is as simple as flattening an equation into one line and should be intuitive for most people, including non-programmers. When it comes to details, conventions used in Python usually take precedence over other conventions. For example, the equation 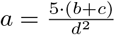 can be written as `a = 5 ∗ (b + c) / d∗∗2`. Here, the power operator is a double asterisk `∗∗` as used in Python. However, the commonly used caret `^` symbol is implemented as a synonym. Parenthesis `(…)` indicate grouping. Arguments to a function are also grouped using parenthesis, e.g. `exp(2)` or `sin(4 + 3)`.

Currently, PyRates does not include a full computer algebra system. By convention, the variable of interest is positioned on the left-hand-side of the equality sign and all other variables and operations on the right-hand-side. First-order differential equations are allowed as an exception: The expression `d/dt ∗ a` is treated as a new variable and can thus be positioned as variable of interest on the left-hand-side as in

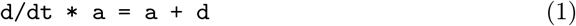

Higher order differential equations must be given as a set of coupled first-order differential equations. For example the equation

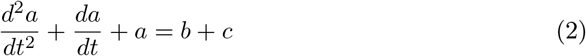

can be reformulated as the following set of two coupled first-order differential equations:

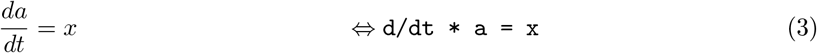

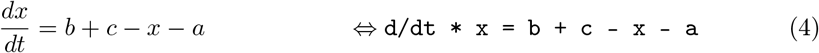

In simulations, this type of equation will be integrated for every time step. The following is an example for equations of a single neural mass in the classic Jansen-Rit model [36], which will be reused in later examples:

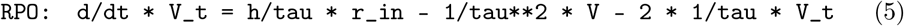

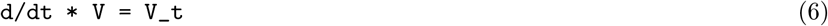

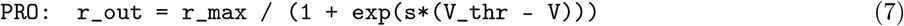

Equation (7) represents the transformation of the population-average membrane potential *V* to an outgoing firing rate *r_out_* via a sigmoidal transformation with slope *s*, maximum firing rate *r_max_* and firing threshold *V_thr_*. This formulation contains a function call to the exponential function via `exp(…)`. Using the preimplemented sigmoid function, equation (7) can be shortened to

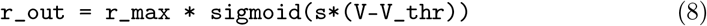

Multiple arguments to a function call are comma separated, e.g. in the sum along the first axis of matrix *A* which would be: `sum(A, 0)`. Using comparison operators as function arguments, it is also possible to encode events, e.g. a spike, when the membrane potential *V* exceeds the threshold *V_thr_*:

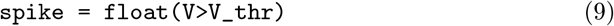

The variable `spike` takes the decimal value `1.0` in case of a spike event and `0.0` otherwise.

The above examples assumed scalar variables, but vectors and higher-dimensional variables are also possible. In particular, indexing is possible via square brackets `[…]` and mostly follows the conventions of *numpy* [40], the *de facto* standard for numerics in Python. Supported indexing methods include single element indexing a[3], slicing `[1:5]`, slicing along multiple axes separated by commas `[0:5,3:7]`, multi-element indexing a[[3], [4]], and slicing via Boolean masks `a[a>5]` for variable a of suitable dimensions. For explanations, please refer to the numpy documentation. A full list of supported mathematical symbols and pre-implemented functions can be found in the supporting information (Tables S1 and S2).

### Components of a network model

In contrast to most other neural simulation frameworks, PyRates treats network models as network graphs rather than matrices. This works well for densely connected graphs, but gives the most computational benefit for sparse networks. Figure 1 gives an overview of the different components that make up a model. A network graph is called a *circuit* and is spanned by *nodes* and *edges*. For a neural population model, one node may correspond to one neural population with the edges encoding coupling between populations. In addition, circuits may be nested arbitrarily within other circuits. Small, self-contained network models can thus easily be reused in larger networks with a clear and intuitive hierarchy. Figure 1 A illustrates this feature with a fictional large-scale circuit which comprises four brain areas and connections between them. Each area may consist of a single node or a more complex *sub-circuit*. Edges between areas are depicted as lines. Figure 1 B zooms into one brain area containing a three-node sub-circuit. This local model corresponds to the three-population Jansen-Rit model [24, 36] with one excitatory (EIN) and one inhibitory interneuron (IIN) population as well as one projecting pyramidal cell (PC) population.

An individual network node consists of *operators*. One operator defines a scope, in which a set of equations and related variables are uniquely defined. It also acts as an isolated computational unit that transforms any number of input variables into one output. Whether an equation belongs to one operator or another decides the order in which equations are evaluated. Equations belonging to the same operator will be evaluated simultaneously, whereas equations in different operators can be evaluated in sequence. As an example, Figure 1 C shows the operator structure of a pyramidal cell population in the Jansen-Rit model. There are two rate-to-potential operators (eqs. (5) and (6)), one for inhibitory synapses (RPOi) and one for excitatory synapses (RPOe). Both RPOs contain identical equations but different values assigned to the parameters. The subsequent potential-to-rate operator (PRO, eq. (7)) sums both synaptic contributions into one membrane potential that is transformed into an outgoing firing rate. In this configuration, the two synaptic contributions are evaluated independently, but possibly in parallel. The equation in the PRO on the other hand will only be evaluated after the synaptic RPOs. The exact order of operators is determined based on the respective input and output variables.

Apart from nodes, edges may also contain coupling operators. An example is shown in Figure 1 D. Each edge propagates information from a source node to a target node. In between, one or more operators can transform the relevant variable, representing coupling dynamics between source and target nodes. This could represent an axon or bundle of axons that propagates firing rates between neural masses. Depending on distance, location or myelination, these axons may behave differently, which is encoded in operators. Note that edges can read any one variable from a source population and can thus be used to represent dramatically different coupling dynamics than those described above.

The described distinction between circuits, nodes, edges and operators is meant to provide an intuitive understanding of a model while giving the user many degrees of freedom in defining custom models.

### Model definition language

PyRates provides multiple interfaces to define a network model in the frontend (see Figure 2). In this section, we will focus on the template interface which is most suitable for users with little programming expertise. All examples are based on the popular Jansen-Rit model [24]. Additionally, we will briefly discuss the implementation of the Montbrió model [38] for completeness.

As described in the previous section, the Jansen-Rit model is a three-population neural mass model whose basic structure is illustrated in Figure 1. The model is formulated in two state-variables: Average membrane potential *V* and average firing rate *r*. Incoming presynaptic firing rates *r_in_* are converted to post-synaptic potentials via the rate-to-potential operator (RPO). In the Jansen-Rit model, this is a second-order linear ordinary differential equation:

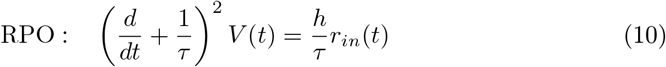

with synaptic gain *h* and lumped time constant *τ*. The population-average membrane potential is then transformed into a mean outgoing firing rate *r_out_* via the potential-to-rate operator (PRO)

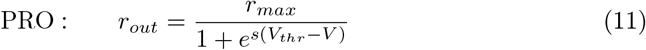

which is an instantaneous logistic function with maximum firing rate *r_max_*, maximum slope *s* and average firing threshold *V_thr_*. The equations above define a neural mass with a single synapse type. Multiple sets of these equations are coupled to form a model with three coupled neural populations. For the two interneuron populations, equation (10) represents synaptic excitation. The pyramidal cell population uses this equation twice with two different parametrizations, representing synaptic excitation and inhibition, respectively. This model can be extended to include more populations or to model multiple cortical columns or areas that interact with each other. For such use-cases PyRates allows for the definition of templates that can be reused and adapted on-the-fly. Templates can be defined using a custom model definition language based on the data serialization standard *YAML* (version 1.2, [41]). The syntax is reduced to the absolute necessities with a focus on readability. The following defines a template for a rate-to-potential operator that contains equation (10):

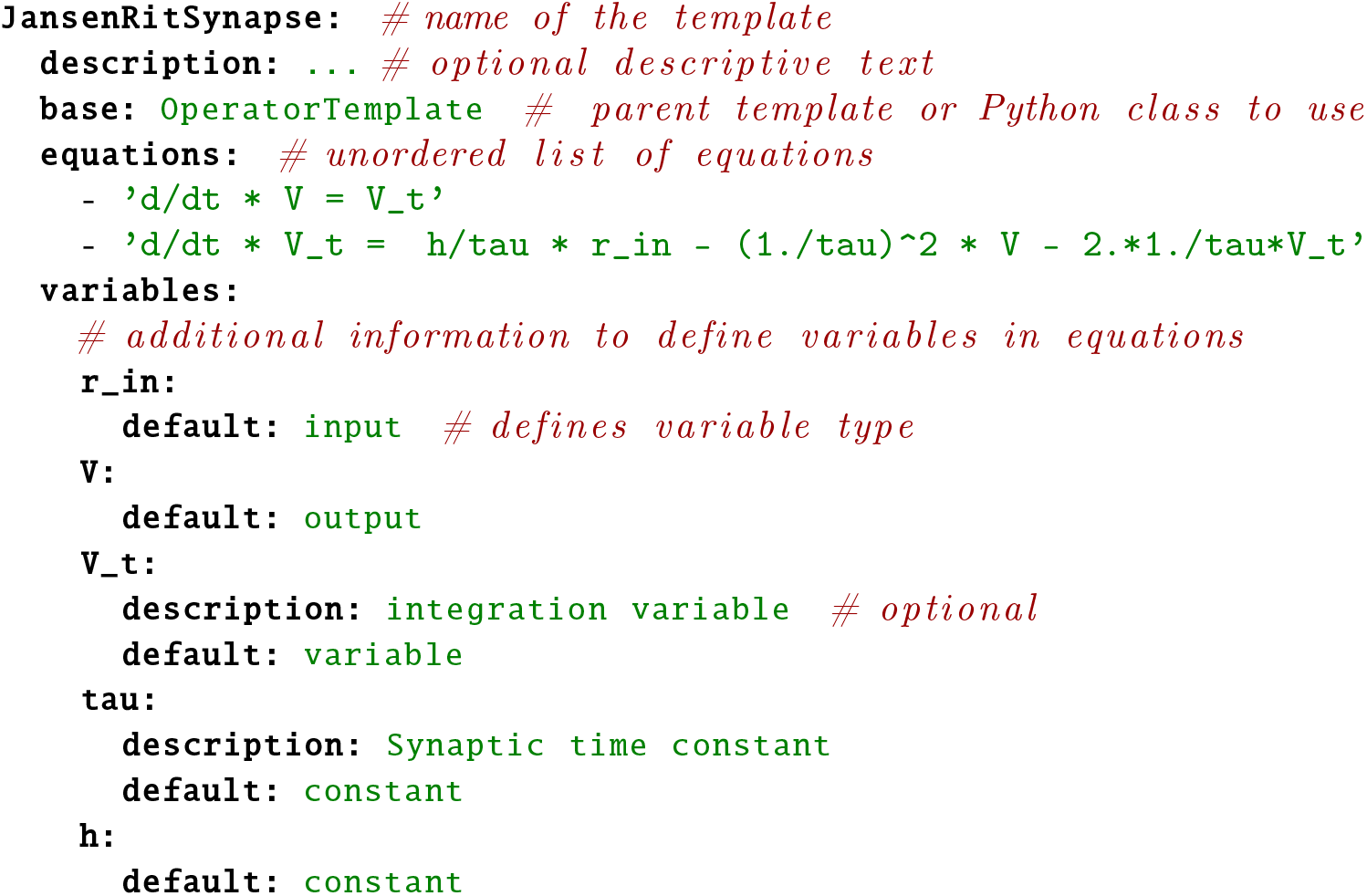

Similar to Python, *YAML* structures information using indentation to improve readability. The base attribute may either refer to the Python class that is used to load the template or a parent template. The *equations* attribute contains an unsorted list of equations, that will be evaluated simultaneously and the *variables* attribute gives additional information regarding the variables defined in *equations*. The only mandatory attribute of variables is *default* which can define the variable type, data type and initial value. Additional attributes can be defined, e.g. a description may help users to understand the template itself or variables in the equations.

For the Jansen-Rit model, it is useful to define sub-templates for excitatory and inhibitory synapses. These share the same equations, but have different values for the constants *τ* and *h* which can be set in sub-templates, e.g. (values based on [36]):

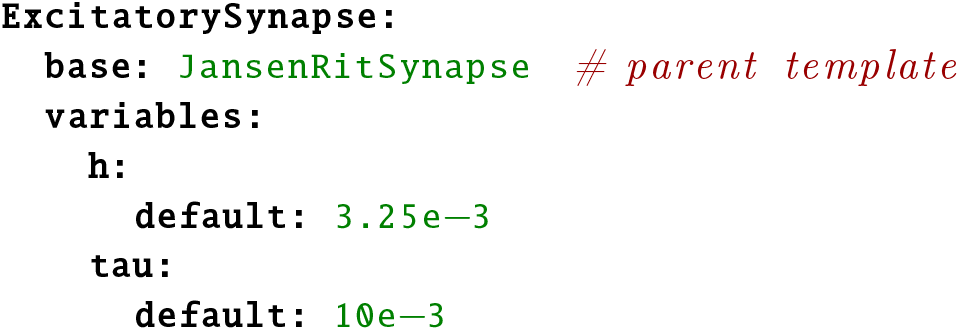

The *JansenRitSynapse* template is reused as *base* template and only the relevant variables are adapted. A single neural mass in the Jansen-Rit model may be implemented as network node with one or more synapse operators and one operator that transforms average membrane potential to average firing rate (PRO, equation (11)/(7)):

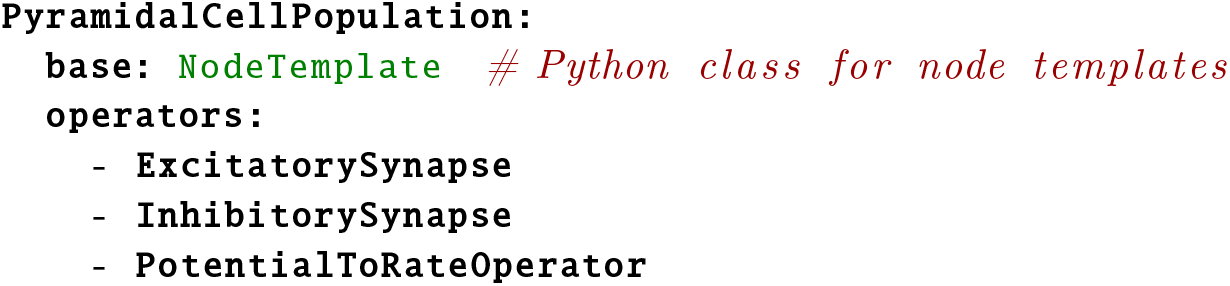

This node template represents the neural population of projecting pyramidal cells as depicted in Figure 1C. The two synapse operators may receive input from other neural masses (or external sources). Equations in these two operators will be evaluated independently and in parallel. The potential-to-rate operator on the other hand sums over the output of the synapse operators and will thus be evaluated after the previous operators. Note that it is also possible to reference an operator template and alter parameter values inside a node template rather than defining separate templates for slight variations.

As described earlier, circuits are used in PyRates to represent one or more nodes and edges between them that encode their coupling. The following circuit template represents the Jansen-Rit model as depicted in Figure 1B:

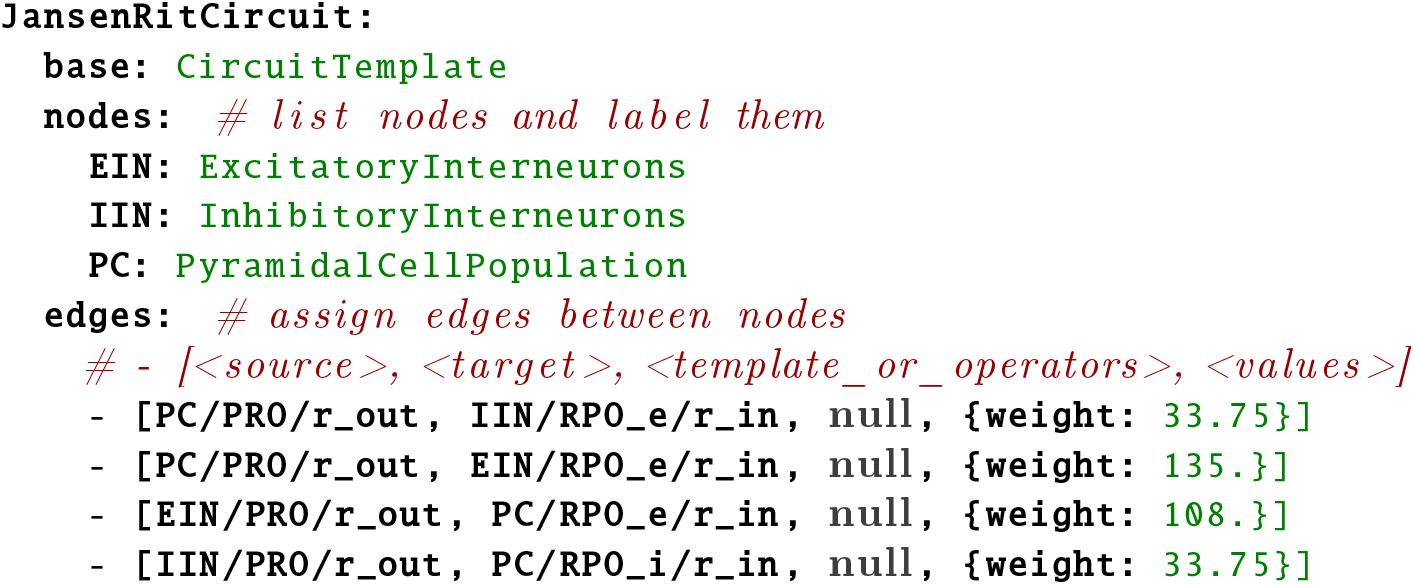

The *nodes* attribute specifies which node templates to use and assigns labels to them. These labels are used in *edges* to define source and target, respectively. Each edge is defined by a list (square brackets) of up to four elements: (1) source specifier, (2) target specifier, (3) template (containing operators), and (4) additional named values or attributes. The format for source and target is `<node_label>/<operator>/<variable>`, i.e. an edge establishes a link to a specific variable in a specific operator within a node. Multiple edges can thus interact with different variables on the same node. Note that the operators were abbreviated here in contrast to the definitions above for brevity. In addition to source and target, it is possible to also include operators inside an edge which do additional transformations that are specific to the coupling between source and target variable. These operators can be defined in a separate edge template that is referred to in the third list entry. In this particular example, this entry is left empty (“null”). The fourth list entry contains named attributes, which are saved on the edge. Two default attributes exist: *weight* scales the output variable of the edge before it is projected to the target and defaults to 1.0; delay determines whether the information passing through the edge is applied instantaneously (i.e. in the next simulation time step) or at a later point in time. By default, no delays are set. Additional attributes may be defined, e.g. to adapt values of operators inside the edge.

In the above example, all edges project the outgoing firing rate *r_out_* from one node to the incoming firing rate *r_in_* of a different node, rescaled by an edge-specific weight. Values of the latter are taken from the original paper by Jansen and Rit [24]. This example with the given values can be used to simulate alpha activity in EEG or MEG.

Jansen and Rit also investigated how more complex components of visual evoked potentials arise from the interaction of two circuits, one representing visual cortex and one prefrontal cortex [24]. In PyRates, circuits can be inserted into other circuits alongside nodes. A template for the two-circuit example from [24] could look like this:

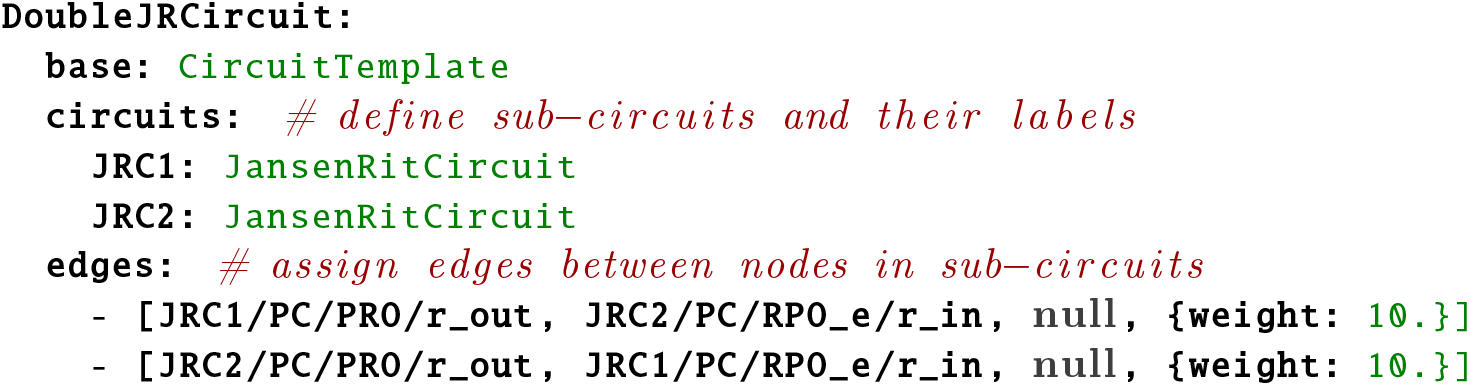

Circuits are added to the template the same way as nodes, the only difference being the attribute name *circuits*. Edges are also defined similarly. Source and target keys start with the assigned sub-circuit label, followed by the label of the population within that circuit and so on.

Besides the *YAML*-based template interface, it is also possible to define models (even templates) from within Python or to implement custom interfaces.

### From model to simulation

All frontend interfaces translate a user-defined model into a set of Python objects which we call the *intermediate representation* (IR, middle layer in Figure 2). This paragraph will give more details on the IR and explain how a simulation can be started and evaluated based on the previously defined model. A model circuit is represented by the `CircuitIR` class, which contains the remainder of the model as a network graph structure using the software package *networkx* [42]. The package is commonly used for graph-based data representation in Python and provides many interfaces to manipulate, analyze and visualize graphs. The `CircuitIR` contains additional convenience methods to plot a network graph or access and manipulate its content. The following lines of code load the `JansenRitCircuit` template that was defined above and transforms the template into a `CircuitIR` instance:

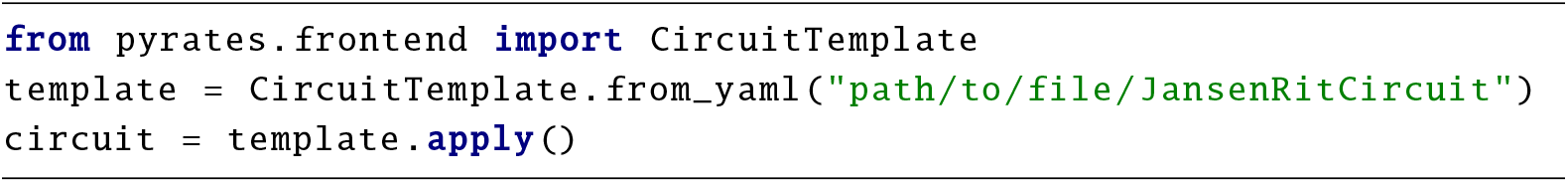

The `apply` method also accepts additional arguments to change parameter values while applying the template. Actual simulations take place in the compute backend (see Figure 2, which again transforms the IR into a more computationally efficient and executable representation. The default backend is based on *tensorflow* [20], which makes use of dataflow graphs to run computations parallelized on one ore more processors (CPUs), graphics cards (GPUs) or across a cluster. The central administrative unit of the backend is the `ComputeGraph` class, which takes care of interpreting equations and optimizing data flow of the model. Optimization mainly consist of summarizing identical sets of (scalar) mathematical operations into more efficient vector operations. The degree of `vectorization` can be specified using the vectorization keyword argument, e.g.:

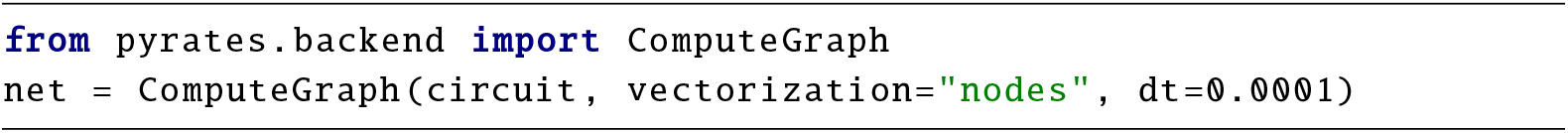

`“none”` indicates that the model should be processed as is; `“nodes”` summarizes identical nodes into one vectorized node; `“full”` also vectorizes identical operators that exist in structurally different nodes, collapsing the entire model into a single node. dt refers to the (integration) time step in seconds used during simulations. Differential equations are integrated using an explicit Euler algorithm which is the most common algorithm used in stochastic network simulations. The unit of dt and the choice of a suitable value depends on time constants defined in the model. Here, we chose a value of 0.1ms,, which is consistent with the numerical integration schemes reported in the literature (e.g. [35, 38]).

A simulation can be executed by calling the run method, e.g.:

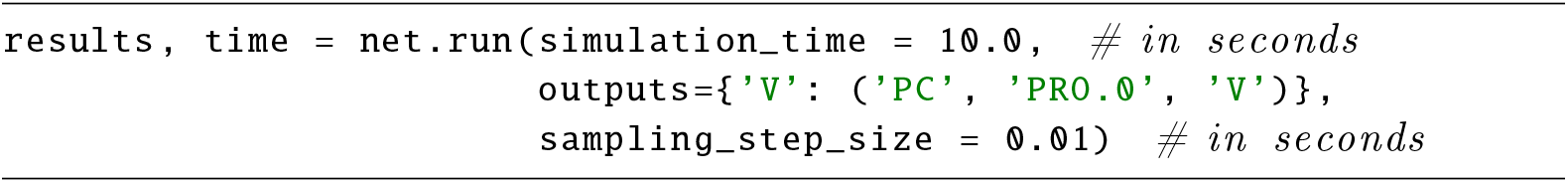

This example defines a total *simulation time* of 10 seconds and specifies that only the membrane voltage from *PC* (pyramidal cell) nodes should be observed. Note that variable histories will only be stored for variables defined as output. All other data is overwritten as soon as possible to save memory. Along this line, a sampling step-size can be defined that determines the distance in time between observation points of the output variable histories. Collected data is formatted as a `DataFrame` from the *pandas* package [43], a powerful data structure for serial data that comes with a lot of convenience methods, e.g. for plotting or statistics. To gain any meaningful results from this implementation of a JRC, it needs to be fed with input in a biologically plausible range. External inputs can be included via *placeholder* variables. To allow for external input being applied pre-synaptically to the excitatory synapse of the pyramidal cells, one would have to modify the `JansenRitSynapse` as follows:

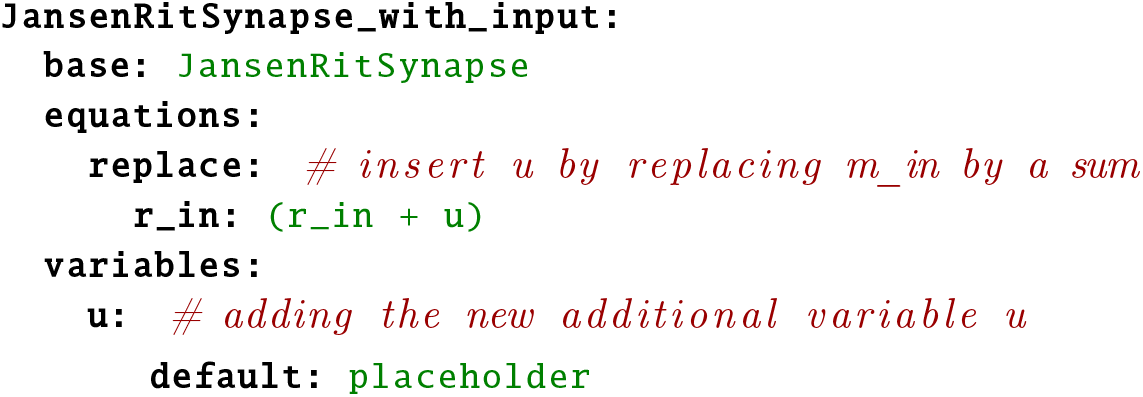

We reused the previously defined `JansenRitSynapse` template and added the variable u as placeholder variable by replacing occurrences of `r_in` by `(r_in + u)` using string replacement. The previously defined equation

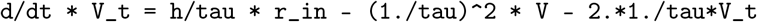

thus turns into

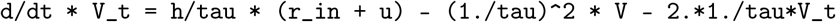

This modification enables the user to apply arbitrary input to the excitatory synapse of the pyramidal cells, using the inputs parameter of the run method:

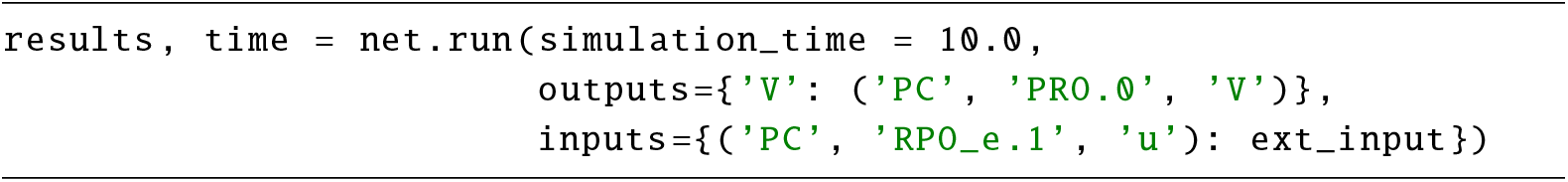

In this example, `external_input` would be an array defining the input value for each simulation step. This subsumes a working implementation of a single Jansen-Rit model that can be used as a base unit to construct models of cortico-cortical networks. By using the above defined *YAML* templates, all simulations described in the next section that are based on Jansen-Rit models can be replicated.

### Implementing the Montbrió model

The neural mass model recently proposed by Montbrió and colleagues is a single-population model that is derived from all-to-all coupled quadratic integrate-and-fire (QIF) neurons [38]. It establishes a mathematically exact correspondence between macroscopic (population level) and microscopic (single cell level) states and equations. The model consists of two coupled differential equations that describe the dynamics of mean membrane potential *V* and mean firing rate *r*:

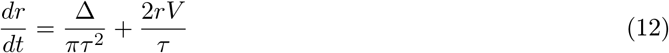

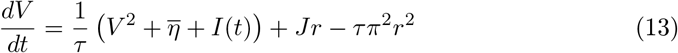

with intrinsic coupling *J* and input current *I*. Δ and 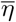 may be interpreted as spread and mean of the distribution of lity excitabi withlevelsin the population. Note that the time constant *τ* was set to 1 and hence omitted in the derivation by Montbrió and colleagues [38]. The following operator template implements these equations in PyRates:

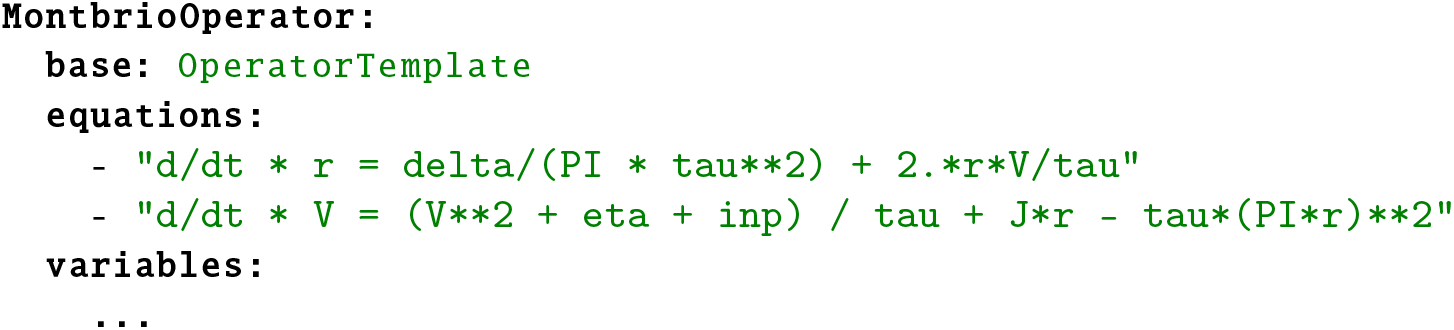

Variable definitions are omitted in the above template for brevity. Since a single population in the Montbrió model is already capable of oscillations, a meaningful network can be set up with a single neural mass as follows:

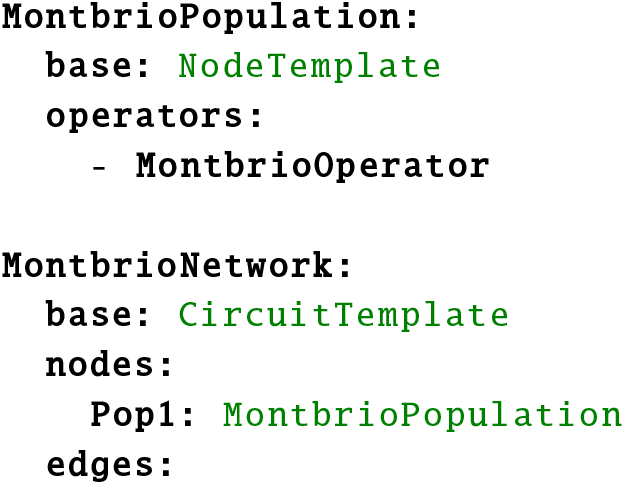

This template can be used to replicate the simulation results presented in the next section that were obtained from the Montbrió model.

### Exploring model parameter spaces

When setting up computational models, it is often important to explore the relationship between model behaviour and model parametrization. PyRates offers a simple but efficient mechanism to run many such simulations on parallel computation hardware. The function `pyrates.utility.grid_search` takes a single model template along with a specification of the parameter grid to sample sets of parameters from. It then constructs multiple model instances with differing parameters and adds them to the same circuit, but without edges between individual instances. All model instances can thus be computed efficiently in parallel on the same parallel hardware instead of executing them consecutively. How many instances can be simulated on a single piece of hardware depends on the memory capacities and number of parallel compute units.

Additionally, PyRates provides an interface for deploying large parameter grid searches across multiple work stations. This allows to split large parameter grids into smaller grids that can be run in parallel on multiple machines.

### Visualization and data analysis

PyRates features built-in functions for quick data analysis and visualization and native support for external libraries due to its commonly used data structures. On the one hand, network graphs are based on *networkx* Graph objects [42]. Hence, the entire toolset of networkx is natively supported, including an interface to the *graphviz* [44] library. Additionally, we provide functions for quick visualization of a network model within PyRates. On the other hand, simulation results are returned as a *pandas.DataFrame* which is a widely adopted data structure for tabular data with powerful built-in data analysis methods [43]. While this data structure allows for an intuitive interface to the *seaborn* plotting library by itself already, we provide a number of visualization functions such as time-series plots, heat maps and polar plots in PyRates as well. Most of those provide direct interfaces to plotting functions from *seaborn* and *MNE-Python*, the latter being an analysis toolbox for EEG and MEG data [45, 46].

Following the principle of modular software design, we prefer to provide interfaces to existing analysis tools, rather than implementing the same functionality in PyRates. In the case of forward-modelled EEG or MEG data, for example, we provide functions that produce Raw, Evoked or Epochs data types expected by *MNE-Python*. A complete list of currently implemented interfaces can be found in the online documentation and more interfaces can be requested or contributed on the public github repository (https://github.com/pyrates-neuroscience/PyRates).

## Results

The aim of this section is to (1) demonstrate that numerical simulations of models implemented in PyRates show the expected results and (2) analyze the computational capabilities and scalability of PyRates on a number of benchmarks. As explained previously, we chose the models proposed by Jansen and Rit and Montbrió and colleagues as exemplary models for these demonstrations. We will replicate the basic model dynamics under extrinsic input as reported in the original publications. To this end, we will compare the relationship between changes in the model parametrization and the model dynamics with the relationship reported in the literature. For this purpose, we will use the grid search functionality of PyRates, allowing to evaluate the model behavior for multiple parametrizations in parallel. Having validated the model implementations in PyRates, we will use the JRC as base model for a number of benchmark simulations. All simulations performed throughout this section use an explicit Euler integration scheme with a simulation step size of 0.1 ms. They have been run on a custom Linux machine with an NVidia Geforce Titan XP GPU with 12GB G-DDR5 graphic memory, a 3.5 GHz Intel Core i7 (4th generation) and 16 GB DDR3 working memory. Note that we provide Python scripts available at https://github.com/pyrates-neuroscience/PyRates/tree/master/documentation which can be used to replicate all of the simulation results reported below.

## Validation of Model Implementations

### Jansen-Rit circuit

The Jansen-Rit circuit is a three-population model that has been shown to be able to produce a variety of steady-state responses [24, 25, 37]. In other words, the JRC has a number of bifurcation parameters that can lead to qualitative changes in the model’s state dynamics. In their original publication, Jansen and Rit delivered random synaptic input between 120 and 320 Hz to the projection cells while changing the scaling of the internal connectivities *C* [24] (reflected by the parameters *C_xy_* in Figure 1B). As visualized in Fig. 3 of [24], the model produced (noisy) sinusoidal oscillations in the alpha band for connectivity scalings *C* = 128 and *C* = 135, thus reflecting a major component of the EEG signal in primary visual cortex. For others, it produced either random noise (*C* = 68 and *C* = 1350) or large-amplitude spiking behavior (*C* = 270 and *C* = 675). We chose to replicate this figure with our implementation of the JRC in PyRates. To this end, we simulated 2 s of JRC behavior for each internal connectivity scaling *C* ∈ {68, 128, 135, 270, 675, 1350} and plotted the average membrane potential of the projection cell population (depicted as *P C* in Figure 1 B). All other model parameters were set according to the parameters chosen in [24]. The results of this procedure are depicted in Figure 3A. While the membrane potential amplitudes were in the same range as reported in [24] in each condition, we re-scaled them for better visualization. As can be seen, they are in line with our expectations, showing random noise for both the highest and the lowest value of *C*, alpha oscillations for *C* = 128 and *C* = 135, and large-amplitude spiking behavior for the remaining conditions. Next to the connectivity scaling, the synaptic time scales *τ* of the JRC are further bifurcation parameters that have been shown to be useful to tune the model to represent different frequency bands of the brains’ EEG signal [25]. As demonstrated by David and Friston, varying these time scales between 1 and 60 ms leads to JRC dynamics that are representative of the delta, theta, alpha, beta and gamma frequency band in the EEG [25]. Due to its practical importance, we chose to replicate this parameter study as well. We systematically varied the excitatory and inhibitory synaptic timescales (*τ_e_* and *τ_i_*) between 1 and 60 ms. For each condition, we adjusted the excitatory and inhibitory synaptic efficacies, such that the product *H_τ_* was kept constant. All other parameters were chosen as reported in [25] for the respective simulation. We then simulated the JRC behavior for 1 min and evaluated the maximum frequency of the power spectral density of the pyramidal cells membrane potential fluctuations. The results of this procedure are visualized in the right panel of Figure 3A. They are in accordance with the results reported in [25], showing response frequencies that range from the delta (1-4 Hz) to the gamma (> 30 Hz) range. Also, they reflect the hyper signal not representative of any EEG signal for too high ratios of 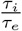. Taken together, we are confident that our implementation of the JRC in PyRates resembles the originally proposed model accurately within the dynamical investigated dynamical regimes. Note, however, that faster synaptic time-constants or extrinsic input fluctuations should be handled carefully. For such cases, it should be considered to reduce the above reported integration step size in order to avoid numerical instabilities.

**Fig 3.**
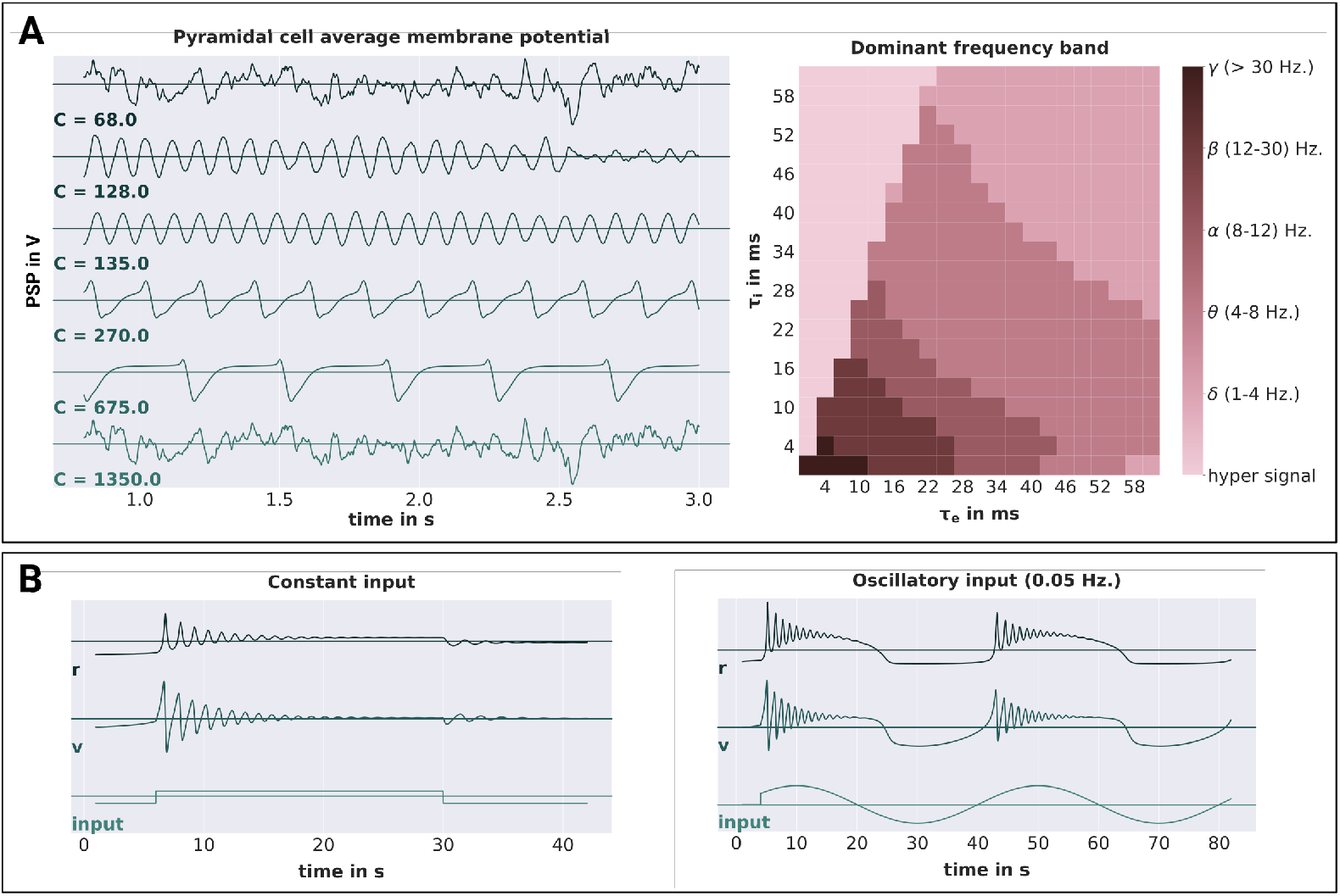
Jansen-Rit and Montbrió model validations. **A** Shows the simulation results obtained from a single Jansen-Rit model. On the left hand side, the average membrane potentials of the pyramidal cell population are depicted for different connectivity scalings C. On the right hand side, the dominant oscillation frequency of the pyramidal cell membrane potentials (evaluated over a simulation period of 60 seconds) is depicted for different synaptic time-scales *τ_e_* and *τ_i_*. The frequencies are categorized into the following bands: *δ* (1-4 Hz), *θ* (4-8 Hz), *α* (8-12 Hz), *β* (12 – 30 Hz), *γ* (> 30 Hz) and h.s. (hyper signal) for signals not representative of any EEG component. **B** Shows the simulation results obtained from a single Montbrió model. The average membrane potentials *υ*, average firing rates *r* and input currents are depicted for constant and oscillatory input on the left and right hand side, respectively.

### Montbrió model

Even though the Montbrió model is only a single-population model, it has been shown to have a rich dynamic profile with oscillatory and even bi-stable regimes [38, 47]. To investigate the response of the model to non-stationary inputs, Montbrió and colleagues initialized the model in a bi-stable dynamic regime and applied (1) constant and (2) sinusoidal extrinsic forcing within a short time-window. For the constant forcing condition, they were able to show that the model responded with distinct damped oscillations to the on- and offset of the forcing, corresponding to two different stable dynamic regimes the model was pushed into (stable focus at onset and stable fixed point at offset). For the oscillatory forcing, on the other hand, the model was pushed from one fixed point (stable node) to the other (stable focus), thereby crossing the bi-stable regime. This behavior can be observed in Fig. 2 in [38] and we chose to replicate it with our implementation of the Montbrió model in PyRates. With all model parameters set to the values reported in [38] for this experiment, we simulated the model’s behavior for 40 s and 80 s for the constant and periodic forcing conditions, respectively. For both conditions, the external forcing strength was chosen as I = 30, while the frequency of the oscillatory forcing was chosen as 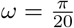. As shown in Figure 3, we were able to replicate the above described model behavior. Constant forcing led to damped oscillatory responses of different frequency and amplitude at on- and offset of the stimulus, whereas oscillatory forcing led to damped oscillatory responses around the peaks of the sinusoidal stimulus. Again, we take this as strong evidence for the correct representation of the Montbrió model by PyRates.

### Benchmarks

Neural simulation studies can differ substantially in the size and structure of the networks they investigate, leading to different computational loads. In this paragraph, we describe how simulation time and memory requirements scale in PyRates as a function of the network size and connectivity. To this end, we simulated the behavior of different JRC networks. Each network consisted of *N* ∈ {2^0^, 2^1^, 2^2^,…, 2^11^} randomly coupled JRCs with a coupling density of p ∈ {0.0, 0.25, 0.5, 0.75, 1.00}. Here, the latter refers to the relative number of pairwise connections between all pairs of JRCs that were established. The behavior of these networks was evaluated for a total of 10 s, leading to an overall number of 10^5^ simulation steps to be performed in each condition (given a step-size of 0.1 ms). To make the benchmark comparable to realistic simulation scenarios, we applied extrinsic input to each JRC and tracked the average membrane potential of every JRC’s projection cell population with a time resolution of 1 ms as output. Thus, the number of input and output operations also scaled with the network size. We assessed the time in seconds and the peak memory in GB needed by PyRates to execute the run method of its backend in each condition. This was done via the Python internal packages *time* and *tracemalloc*, respectively. Thus, all results reported here refer to the mere numerical simulation, excluding the model initiation. To account for random fluctuations due to background processes, we chose to report the averages of simulation time and peak memory usage over *N_R_* = 10 repetitions of each condition. To provide an intuition of these fluctuations, we calculated the average variation of the simulation time and peak memory usage over conditions as 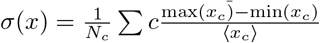, with *x* being either the simulation time or the peak memory usage, *c* being the condition index and 〈*x*〉 representing the expectation of x. We found the simulation time and peak memory consumption to show average variations of 2.9260*s* and 0.0007*M B*, respectively, which is in both cases orders of magnitude smaller than the variations of the mean simulation time and memory consumption we found across conditions. The average simulation times are visualized in Figure 4A. They demonstrate the effectiveness of PyRates’ backend in parallelizing network computations, since the simulation time practically did not scale with the network size and coupling density for the largest part of the conditions. Furthermore, they demonstrate the current upper limit of the parallelization, since for networks of *N* ≥ 2048 JRCs, the simulation time scales linearly with increases in the coupling density *p*. However, we expect new hardware developments in the GPU sector to raise this upper limit. Importantly, the efficiency of our tensorflow-based backend does not rely entirely on the availability of strong GPUs. In Figure 4B, we show the ratio between the benchmark simulations performed on the GPU vs. the same benchmarks run on the CPU. We found that running simulations on the CPU can even be faster for small- to mid-sized as well as all uncoupled networks within the investigated network size range. When it comes to networks with *N* ≥ 256 JRCs, the advantage of GPUs over CPUs becomes substantial, though, with simulations that were up to 50 times faster on the GPU.

**Fig 4.**
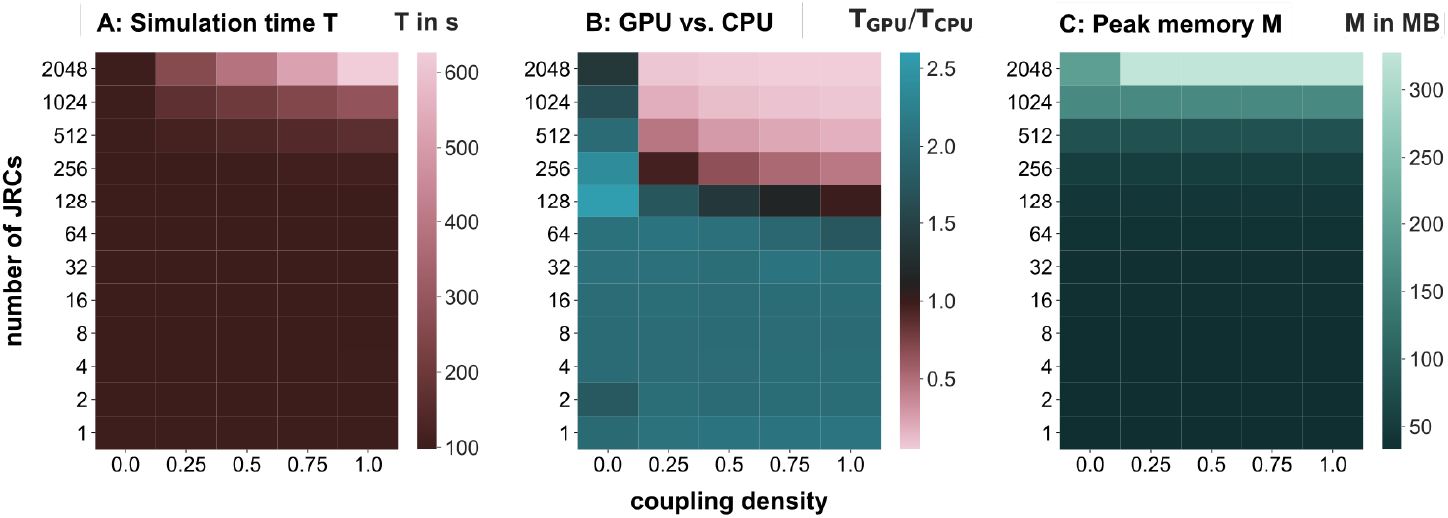
PyRates benchmarks. Benchmark results for 1 min simulations run in PyRates with a simulation step-size of 0.1 ms. Simulations were performed for networks with different numbers of Jansen-Rit circuits (JRCs) and differently dense coupling between the JRCs (connection probability). **A** shows the average simulation time on the GPU, **B** shows the ratio between the average simulation time on the GPU and the CPU, and C shows the average peak memory consumption independent of the device.

Regarding memory requirements, Figure 4C shows that the peak memory consumption mostly scaled with the size of the network. This reflects (1) the independence between the size of the simulation output and the coupling density p and (2) that memory requirements are mostly determined by the size of the simulation output and to a lesser extent by the size of the underlying graph representation of the network.

It is important to note that the benchmark results reported here also hold for parameter sweeps performed via the grid search functions of PyRates, as they exploit exactly the same parallelization mechanisms. If, for example, a user would analyze the behavior of a single JRC for different parametrizations of the model (as we did in the first part of this section), the dependency of the simulation time on the number of different model parametrizations would be expressed by column 1 of Figure 4A. This makes PyRates especially powerful for the exploration of model parameter spaces.

## Discussion

In this work we presented PyRates, a novel Python framework for designing neural models and numerical simulations of their dynamic behavior. We introduced the frontend, including its user interfaces, structure and mathematical syntax, as well as the *tensorflow*-based backend. For validation purposes, we implemented the neural population models proposed by Jansen an Rit and Montbrió and colleagues in PyRates and successfully replicated their key dynamic features reported in the literature [24, 38]. This supports that both the model configurations produced by our frontend and their translation into tensorflow compute graphs by our backend are accurate. Additionally, we tested the computational power of our backend on a number of different benchmarks. Those benchmarks consisted of simulations of JRC networks that differed in the number of their nodes and edges. On our computing hardware, these benchmark simulations revealed simulation times that practically did not scale with the network size and coupling density for small- to medium-sized networks, due to the efficient parallelization. We showed how model parameter sweeps benefit from this parallelization and for which problem sizes to use the CPU or GPU to run simulations on. From these results, we conclude that PyRates is a powerful simulation framework that allows for highly efficient brain network simulations. The main questions we will address in the following discussion are why PyRates is a valuable addition to established neural simulation software and in which cases scientists can benefit from using it.

### PyRates in the context of existing neural simulation frameworks

Within the domain of neural simulation frameworks, PyRates belongs to the family of graph-based neural simulators. In both its frontend and backend, it represents a neural model as a network of nodes connected by edges. Inherently, PyRates makes no assumptions about the spatial scale of nodes and edges in its networks, thus rendering it feasible for neural networks of any type. Additionally, PyRates allows for merging and hierarchical organization of neural networks by building graphs from sub-graphs. Hence, our tool can also be used to build multi-scale models, *e.g*. a macroscopic network of connected neural populations, with some populations of interest being represented by sub-networks of single neurons. This being said, PyRates has only been systematically tested on rate-based population models, yet. These differ qualitatively from spiking neuron models in their output variable, which is continuous for rate-based models but discrete for spiking neuron models. While it is in principle possible to implement such discrete spiking mechanisms, the compute engine is not optimized for it, since it projects output variables at each time-step to their targets in the network. This means that the projection operation will be performed regardless of whether a spike is produced or not, leading to considerable increases in computation time for large, densely connected single cell networks. Hence, when dealing with neuroscientific questions that implicate the use of spiking neuron models, we currently recommend to use simulation tools such as Nengo [13], NEST [14], ANNarchy [15], Brian [16] or NEURON [17]. Such questions may involve problems where specific spike-timings have a non-negligible influence, where dendritic tree architectures are important or, generally, where the variable of interest looses its meaning when averaged over time or over many neurons.

Of course, all of the above listed tools can be applied in other scenarios as well, even for macroscopic neural network simulations. However, if the variable of interest in a given model can be expressed as an average over many cells and single cell dynamics can be neglected, mean-field approaches such as the neural population models used throughout this article will be considerably faster and thus allow for the investigation of larger networks and parameter spaces. In general, most frameworks that feature generic code generation should allow to implement such models. From the above mentioned tools, Brian and ANNarchy belong to that category. Brian is strictly aimed at spike-based simulations and thus not optimized for continuous output variables like firing rates, whereas ANNarchy provides features for spike- and rate-based neural simulations. Nonetheless, it is designed for single-cell network simulations, so most of the templates it provides for neurons or populations are not necessarily applicable to mean-field models. Other simulation frameworks that provide explicit mean-field modeling mechanisms include TVB [48], DCM [12], DiPDE [49] and MIIND [50]. Among these, the latter two focus strongly on so-called population density techniques, which can describe the full voltage probability distribution of a population of neurons, instead of its mere mean. Both DiPDE and MIIND focus on the leaky integrate-and-fire neuron as the underlying neuron model to derive the voltage probability distribution from. The advantage of this technique is the more direct and precise relationship between the single cell activity and the population level as compared to mean-field approaches. However, this advantage is payed for by higher computational demands, since a discretized probability distribution is computed at each simulation step instead of a mere point-estimate (i.e. the mean). TVB and DCM, on the other hand, focus on the same mathematical group of neurodynamic models as currently implemented in PyRates, i.e. neural population models. The focus of TVB lies in the simulation of large-scale brain networks via established, preferably homogeneous local population models and DCM is explicitly designed to infer parameters of a fixed set of pre-implemented models based on a given measure of brain activity. While being the optimal choice for their respective use-cases, both tools lack functionalities that help implementing custom models.

We consider the core strengths of PyRates to be its highly generic model definition (comparable to a pure code generation approach) and its tensorflow-based backend. The former distinguishes PyRates from other simulation frameworks, since it allows to customize every part of a neural network as long as a network structure with nodes and edges defined by mathematical operators is maintained. Every single computation that is performed in a PyRates simulation and every variable that it performs on is defined in the frontend and can be accessed and edited by the user. This allows, for example, to add custom synapse types, plasticity mechanisms, complex somatic integration mechanisms or even axonal cable properties. In addition, edges can access and connect all variables existing on pre- and post-node, thus enabling the implementation of projections or plasticity mechanisms that depend on population variables other than firing rates. This generic approach makes PyRates particularly valuable for neuroscientists interested in developing novel neural models or extending existing ones. A notion of care should be added here, however. The degrees of freedom we provide for setting up models and simulations in PyRates imply that we do not provide safeguards for questionable model definitions. Except for their syntactical correctness, model equations and their hierarchical relationships will not be questioned further by PyRates. Also, inputs and outputs to the model will be added exactly as defined by the user. In other words, while PyRates does provide a considerable number of convenience functions to quickly set up and simulate large neural networks, it still requires the users to be aware of potential numerical issues they could run into, if the model or simulation would not be set up correctly. Typical pitfalls include numerical overflows if variables become to large or small for the chosen data type, simulation step sizes that were chosen too large for the internal timescales of a given model, and random variables which are sampled at each simulation step without taking into account the dependency between sampling frequency and simulation step size. We tested numerical solvers providing adaptive time steps as an alternative to the Euler algorithm to handle the problem of choosing an appropriate integration step size. However, we found those algorithms to be unsuited for network simulations in PyRates, since handling asynchronicity between network nodes created significant computational overhead. Regarding PyRates’ second core strength, its backend, we have demonstrated its computational power in various scenarios. It provides efficient representations of large neural networks that are highly optimized for parallel execution on CPUs and GPUs. Parallel execution of network simulations are particularly efficient when its nodes and edges are similar in their mathematical operators, since those similarities are exploited by the automatic vectorization mechanisms of PyRates. In turn, this means that the effectiveness of the parallelization scales negatively with the relative amount of heterogeneity or sequentiality of the network. Networks that consist of highly diverse neural units governed by many, hierarchically dependent operators will show considerably longer simulation times than networks with very similar elements and a flat operator hierarchy. Thus, PyRates is particularly suited for simulating large, homogeneous networks or conducting parameter studies on small- to medium sized networks. For the latter, we provide functionalities that take a network and a number of different parameter sets of that network as input, organizes the different parametrizations of the network in a new network and simulates their behavior in a single simulation. This way, parameter searches on shared-memory systems are substantially sped up. Furthermore, we provide a cluster distribution mechanism that makes this feature available for distributed simulations on large compute clusters.

### Integrating PyRates into neuroscientific work-flows

Neural population models such as the Jansen-Rit model [24] were originally conceived to understand or predict physical measures of brain activity such as LFPs, EEG/MEG or BOLD-fMRI. Modern neuroscientific workflows, however, go beyond mere forward simulations of brain activity. For example, The Virtual Brain [48] allows the use of structural (including diffusion-weighted) MRI scans to specify 3-dimensional structure and connectivity of a network design. Dynamic Causal Modeling [12] on the other hand can make use of measured brain activity to infer model parameters (e.g. connectivity constants) that best fit the given data. Both approaches have in common, that brain network models are adapted to individual subjects based on measured data. PyRates integrates well with this concept for two reasons: (1) It is designed to provide an easy-to-use interface to construct and adapt network models with more flexibility than comparable tools. (2) Due to its modular software structure, PyRates can easily be extended to interface with existing tools. While the intermediate representation serves as a standard interface, front- and backend can be exchanged to integrate with other software. For example, PyRates could be extended with a frontend that makes use of structural MRI data via tools provided by TVB. At the same time, the current backend could be replaced with a code generator that produces region-specific models compatible with TVB’s node model interface. PyRates can thus provide more elaborate, task- or region-specific models and integrated with tools like TVB or DCM that combine simulations with measured data.

Currently, PyRates already provides a number of useful interfaces to tools that can be used for setting up models or subsequent analyses. Two of those interfaces come with the graph representations PyRates uses for networks. As mentioned before, every PyRates network is translated into a *tensorflow* graph. This enables the usage of every tensorflow function that could come in handy for setting up a model in PyRates, be it mathematical functions like *sine* or *max*, variable manipulation methods like *reshape* or *squeeze* or higher-level functions like error measurements or learning-rate decays. For the future, we also plan to provide interfaces to *tensorflow’s* model training features, thus allowing to optimize parameters of neural models via gradient-descent based algorithms [20]. Since the intermediate representation fully builds on networkx graphs, the networkx API can be used to create, modify, analyze or visualize models. This includes interoperability with explicit graph visualization tools like Graphviz [43] or Cytoscape [51] that contain more elaborate features for visualizing complex biological networks. For the processing, analysis and visualization of simulation results, we provide a number of tools that mostly wrap *MNE-Python* [45, 46] and seaborn [52] functions. For extended use of *MNE-Python*, we also provide a wrapper that allows to translate every output of a PyRates simulation into an *MNE-Python* object. This is particularly useful for forward simulations of EEG/MEG data, since *MNE-Python* comes with an extensive range of methods for the processing, analysis and visualization of such data. Finally, PyRates can also be used in combination with pygpc, a generalized polynomial chaos (GPC) toolbox for uncertainty quantification and sensitivity analysis publicly available under https://github.com/konstantinweise/pygpc. Via this interface it is possible to define a model plus a set of model parameters, including their respective uncertainties, and estimate how sensitive the model behavior is to changes in these parameters. It is important to notice however, that the GPC cannot replace a proper bifurcation analysis and should currently only be used for parameter ranges where no bifurcations or multi-stabilities occur. In summary, PyRates is readily integrated into complex neuroscientific workflows as a tool for bottom-up neural simulations. To this end, it provides interfaces to other Python tools that have been specifically designed to manage other parts of such workflows (e.g. data processing, visualization,…). More interfaces can easily be implemented due to the modular structure of the framework. This is further aided by the widely used data structures PyRates is build on, like YAML-based configuration files, networkx graphs or pandas DataFrames. PyRates can thus be included as one independent component of larger neuroscientific workflows.

## Acknowledgments

Richard Gast has been supported by the Max Planck Society and is currently funded by the Studienstiftung des Deutschen Volkes. Daniel Rose is supported by the International Max Planck Research School NeuroCom. Nikolaus Weiskopf is supported by the European Research Council under the European Union’s Seventh Framework Programme (FP7/2007-2013) / ERC grant agreement no. 616905, the BMBF (01EW1711A & B) in the framework of ERA-NET NEURON, the European Union’s Horizon 2020 research and innovation programme under the grant agreement No 681094.

## Supporting information

The mathematical syntax in PyRates is based on conventions used in the Python community. Table S1 summarizes the most common operations. Variable and function names consist of latin letters, underscores and numbers but must start with a letter or underscore. Indexing of higher-dimensional variables follows the numpy convention [40].

Table S2 shows preimplemented mathematical functions that are exposed in the mathematical syntax. Note that tensorflow may support more functions than PyRates exposes to the user. It is, however, straightforward to extend the list of exposed functions as needed. For a complete list of functions supported by tensorflow, please refer to the tensorflow API documentation [20].

**Table S1.** Overview of mathematical syntax.

| Syntax | Description | Syntax | Description |
| --- | --- | --- | --- |
| **basic operations** |  | **reserved constants** |  |
| a+b | addition | E, e | Euler's number |
| a-b | subtraction | PI | pi ($\pi$, Archimedes' constant) |
| a*b | multiplication | Parentheses |  |
| a/b | division | (...) | prioritize operation, also function calls |
| a%b | modulo | a[...] | indexing |
| -a | negation | **indexing (following numpy standard)** |  |
| a**b, a^b | exponent | a[b] | access element b of array a |
| **matrix operations** |  | a[b:c] | slice: access elements from b to c, excluding c. b or c may be left out. |
| a@b | matrix multiplication | a[b,c] | access element on index a_{b,c} |
| a.T | matrix transpose | a[[b], [c]] | multi-element indexing |
| a.I | matrix inverse | a[a>b] | boolean mask: access all elements for which the condition is true (the length of the resulting array must be the same as the length of the respective axis) |
| **relations** |  |  |  |
| a>b | greater |  |  |
| a<b | less |  |  |
| a==b | equal |  |  |
| a!=b | not equal |  |  |
| a>=b | greater or equal |  |  |
| a<=b | less or equal |  |  |

The syntax follows conventions defined by the Python programming language and the numerics package numpy with few additions. Mathematical functions are mapped on functions provided by the compute engine tensorflow.

**Table S2.**
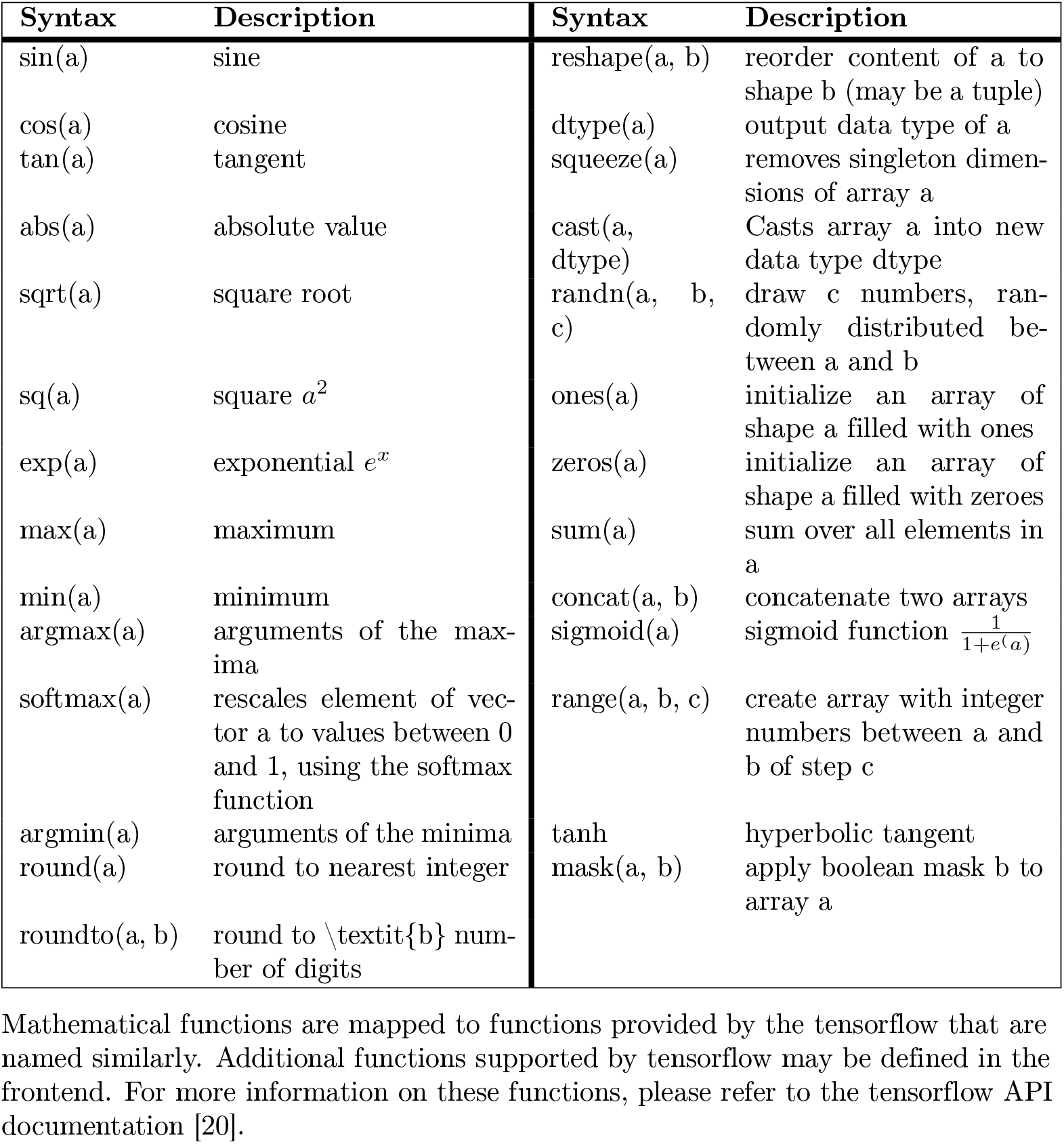
Overview of preimplemented mathematical functions.

| Syntax | Description | Syntax | Description |
| --- | --- | --- | --- |
| sin(a) | sine | reshape(a, b) | reorder content of a to shape b (may be a tuple) |
| cos(a) | cosine | dtype(a) | output data type of a |
| tan(a) | tangent | squeeze(a) | removes singleton dimensions of array a |
| abs(a) | absolute value | cast(a, dtype) | Casts array a into new data type dtype |
| sqrt(a) | square root | randn(a, b, c) | draw c numbers, randomly distributed between a and b |
| sq(a) | square $a^2$ | ones(a) | initialize an array of shape a filled with ones |
| exp(a) | exponential $e^x$ | zeros(a) | initialize an array of shape a filled with zeroes |
| max(a) | maximum | sum(a) | sum over all elements in a |
| min(a) | minimum | concat(a, b) | concatenate two arrays |
| argmax(a) | arguments of the maxima | sigmoid(a) | sigmoid function $\frac{1}{1+e^{(a)}}$ |
| softmax(a) | rescales element of vector a to values between 0 and 1, using the softmax function | range(a, b, c) | create array with integer numbers between a and b of step c |
| argmin(a) | arguments of the minima | tanh | hyperbolic tangent |
| round(a) | round to nearest integer | mask(a, b) | apply boolean mask b to array a |
| roundto(a, b) | round to \textit{b} number of digits |  |  |

Mathematical functions are mapped to functions provided by the tensorflow that are named similarly. Additional functions supported by tensorflow may be defined in the frontend. For more information on these functions, please refer to the tensorflow API documentation [20].

